# Molecular Insights into *ANPEP* in Gastric Adenocarcinoma

**DOI:** 10.1101/2025.08.28.672866

**Authors:** Taíssa Maíra Thomaz Araújo, Bianca de Fátima dos Reis Rodrigues, Jessica Manoelli Costa da Silva, Myrth Soares do Nascimento Remígio, Fabiano Cordeiro Moreira, Samir Mansour Moares Casseb, Williams Fernandes Barra, Geraldo Ishak, Ana Karyssa Mendes Anaissi, Leandro Magalhães, Amanda Vidal, Ronald Matheus da Silva Mourão, Eliel Barbosa Teixeira, Diego Pereira, Valéria Cristiane Santos da Silva, Daniel de Souza Avelar, Rubem Ferreira Silva, Ândrea Kely Ribeiro dos Santos, Livia Erika Carlos Marques, Rommel Rodriguez Burbano, Paulo Pimentel de Assumpção

## Abstract

*Alanyl aminopeptidase* (*ANPEP*) has been implicated in various cancers, but its specific role in gastric adenocarcinoma (GC) remains incompletely understood. This study analyzed *ANPEP* gene expression in gastric cancer (GC), peritumoral tissue (PTT), metaplasia (M), and normal tissue (N). Total RNA was extracted, libraries were prepared and sequenced on the Illumina NextSeq 500. Data was processed using the nf-core/rnaseq pipeline. Transcript quantifications were imported with tximport and normalized using DESeq2. Differential expression (|log_2_FC| >2; adj. p < 0.05) and Kruskal–Wallis tests identified key genes. *ANPEP* was significantly upregulated in GC, PTT, and M compared to normal tissue (p < 0.01), suggesting its involvement in early mucosal transformation and malignant progression. Heatmap analysis revealed upregulation of genes related to immune function and oxidative stress, indicating an immunosuppressive and apoptosis-resistant tumor microenvironment. Correlation analyses identified strong positive associations between *ANPEP* and genes involved in cytoskeletal remodeling, immune modulation, and metabolic regulation, suggesting that *ANPEP* supports both the invasive potential of tumor cells and the establishment of an immunosuppressive niche. These findings position *ANPEP* as a promising biomarker for early detection and a candidate for targeted therapies.

## INTRODUCTION

Gastric cancer (GC) remains a significant global health challenge, ranking among the leading causes of cancer-related mortality worldwide (SUNG *et al*., 2021). Despite advances in early detection and therapeutic strategies, the molecular mechanisms underlying GC progression and metastasis are still incompletely understood, necessitating the identification of novel biomarkers and therapeutic targets (MATSUOKA *et al*., 2024). Among the genes implicated in GC pathogenesis, *ANPEP*/CD13 (*Aminopeptidase N*, also known as *CD13*) has emerged as a promising candidate due to its multifaceted roles in tumor biology (WICKSTRÖM *et al*., 2011).

*ANPEP* encodes a zinc-dependent metalloprotease expressed on the surface of various cell types, including epithelial and endothelial cells, where it regulates processes such as cell migration, invasion, and angiogenesis (LENDECKEL *et al*., 2023). In the context of cancer, *ANPEP* is frequently overexpressed in tumor tissues and is associated with aggressive phenotypes, including enhanced metastatic potential and resistance to therapy (XIU *et al.*, 2022; HU *et al*., 2020). Recent studies have highlighted its significance in gastrointestinal malignancies, particularly GC, where *ANPEP* overexpression correlates with poor prognosis and advanced disease stages (ZHANG *et al*., 2023).

For instance, XIU et al. (2022) demonstrated that *ANPEP* promotes tumor cell invasion in GC by modulating the extracellular matrix and facilitating epithelial-to-mesenchymal transition (EMT), a critical step in metastasis.

The interplay between *ANPEP* and signaling pathways, such as Wnt/B-catenin and PI3K/Akt, further underscores its relevance as a therapeutic target (GUO *et al*., 2019; Wang et al., 2015). Studies have identified *CD13* as a CSC-specific membrane marker, including for hepatocellular carcinoma (HCC), and cholangiocarcinoma (CUI *et al.*, 2018; LAUKKANEN *et al*., 2016). From a functional point of view, *CD13* is known to protect CSC from apoptosis (KESAVARDHANA *et al.*, 2018). Pharmacologically inhibiting CD13 enzymatic activity, the antitumor effects of chemotherapeutic agents, e.g., 5-fluorouracil, can be enhanced (XIU *et al*., 2022; SUN *et al*., 2015).

HOFT *et al*. (2024) reported that identification of a metaplastic cell expressing the cancer-associated biomarker *ANPEP*, present in Hp-induced gastritis and autoimmune atrophic gastritis, indicates the carcinogenic capacity of both diseases. This finding may guide early detection and risk stratification for patients with chronic gastritis.

This study aims to consolidate current knowledge on the role of *ANPEP* in gastric cancer, focusing on its molecular mechanisms, clinical implications, and potential as a therapeutic target. By synthesizing findings from recent studies, we seek to provide insights into how *ANPEP* contributes to GC progression and highlight avenues for future research and clinical translation.

## MATERIAL AND METHODS

### Ethical Approval and Patient Selection

This study was approved by the Research Ethics Committee (CEP) under protocol number CAAE 47580121.9.0000.5634. All enrolled participants provided written informed consent in accordance with the Declaration of Helsinki. Samples of tumor tissue (286) and adjacent non-tumor tissue (61)were collected from patients with histologically confirmed gastric adenocarcinoma. In addition to these, samples from intestinal metaplasia (20)and non-cancerous gastric tissues (27) were also included for comparative purposes.

## RNA Extraction and Sequencing

Total RNA was extracted using the TRIzol® reagent (Invitrogen),following the manufacturer”s protocol. RNA quality and integrity were verified prior to library preparation. Libraries were constructed using the TruSeq Stranded Total RNA kit (Illumina®), and sequencing was performed in paired-end mode on the Illumina NextSeq 500® platform.

### Dry Lab Pipeline

#### nf-core/rnaseq

RNA sequencing data were processed using the nf-core/rnaseq pipeline(Ewels *et al*., 2020)., a community-curated workflow that ensures reproducibility and consistency in RNA-seq analyses. The pipeline performs quality control (QC), adapter trimming, and read alignment or pseudo-alignment using STARand Salmon, respectively. The final outputs include a gene-level expression matrix.

### Import and Normalization of Quantified Reads

Transcript-level quantifications generated by Salmon were imported into the R statistical environment (R Core Team, 2018). using the tximport package (Love *et al*., 2017). This step enables aggregation of transcript abundances to the gene level while correcting for transcript length and library size. The resulting count matrices were normalized using the DESeq2 package, which estimates size factors to account for differences in sequencing depth and sample composition. Normalized counts were extracted for downstream analyses.

### Differential Expression Analysis

Differential gene expression analysis was performed between tumor and adjacent non-tumor tissues using DESeq2. Genes were considered differentially expressed (DE) if they exhibited an absolute log_2_ fold-change greater than 2 and an adjusted p-value (Benjamini–Hochberg correction) below 0.05. Additionally, the Kruskal–Wallis test was applied (p < 0.01) to identify further gene expression differences across clinical or microbiological subgroups, providing a non-parametric validation of DE patterns.

### Correlation and Integrative Analysis

To investigate associations among gene expression, clinical features, and microbial composition, Spearman”s rank correlation was calculated between normalized gene expression values or clinical metadata. This integrative approach enabled the identification of biologically meaningful interactions within the tumor microenvironment.

### Clustering and Data Visualization

For unsupervised clustering and visualization, Kendall rank correlation matrices were generated and used for hierarchical clustering of the DE gene set. Clustered heatmaps were constructed using the ComplexHeatmap package in R, allowing the identification of co-regulated gene modules and biologically informative sample groupings reflective of tumor subtype, clinical condition, or microbial context.

## RESULTS

### Expression of *ANPEP*

Analysis of *ANPEP* gene expression was performed across a cohort of 394 samples, including gastric adenocarcinoma (GC), peritumoral tissue (PTT), metaplasia (M), and normal tissue samples (N). *ANPEP* exhibited variable expression across sample types, with significantly higher expression in GC, PTT, and M compared to normal tissues (p < 0.01, Kruskal-Wallis test), suggesting that *ANPEP* may be upregulated early during gastric mucosal transformation and remain elevated through to malignant progression (Figure 1).

**Figure 1.**
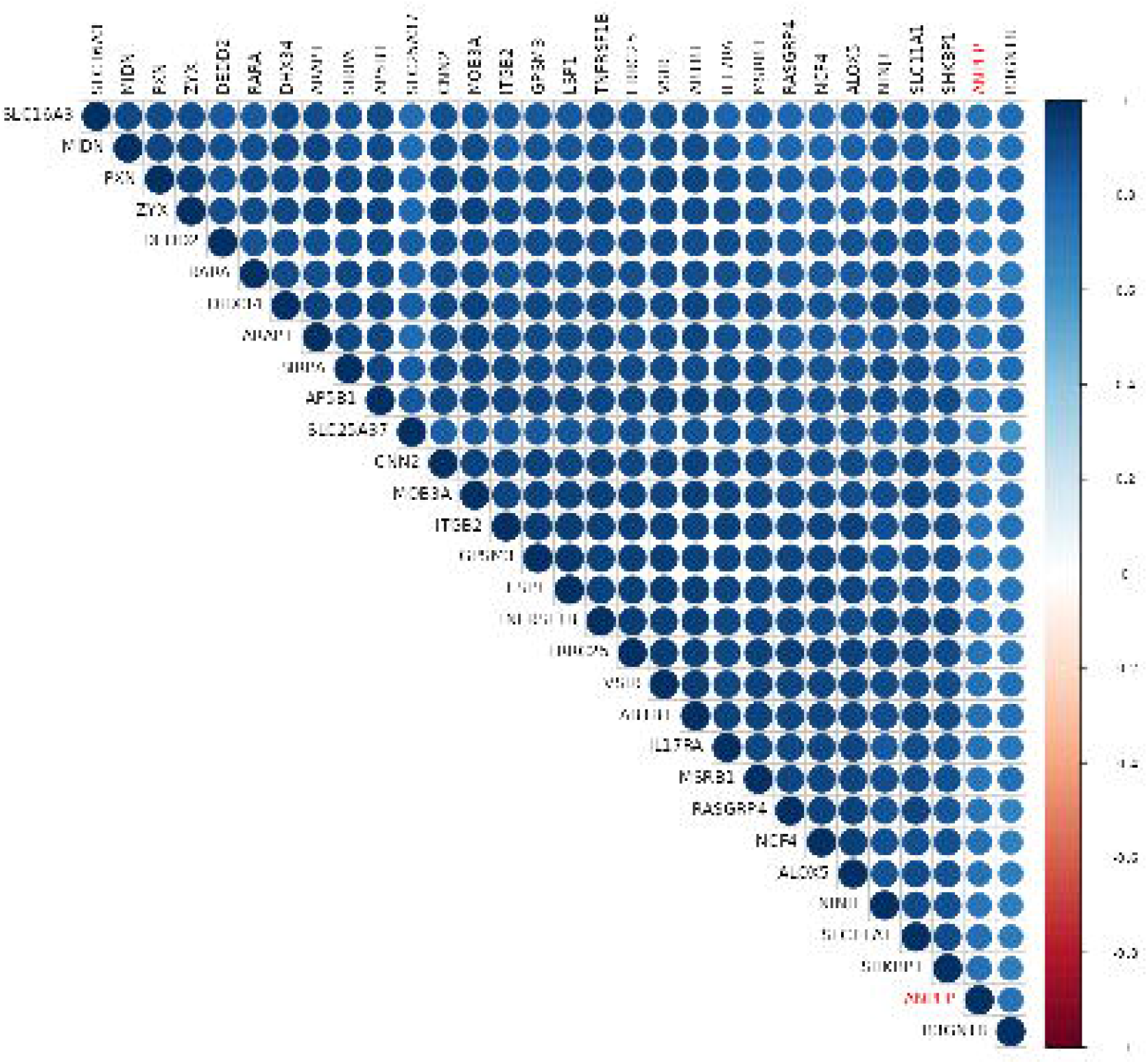
Boxplot of ANPEP expression in gastric adenocarcinoma (GC), peritumoral tissue (PTT), metaplasia (M), and normal tissue samples (N). p < 0.01, Kruskal-Wallis test.

*ANPEP* gene expression does not significantly correlate with clinicopathological variables in gastric adenocarcinoma, including Lauren classification (histological type) – p = 0.592; Neoadjuvant treatment status – p = 0.204; EBV and *H. pylori* infection – p > 0.3; Clinical staging (pT, pN, pM, overall stage) – p > 0.4; Sex, tumor differentiation, and topography – p > 0.05.

### Heatmap

Analysis of the gene expression heatmap revealed a cluster of genes significantly overexpressed in gastric cancer tissues (Figure 2)., many of which are implicated in key oncogenic processes. Notably, *TNFRSF1B, VSIR*, and *SIRPA* are immune-related genes involved in immune evasion, T cell suppression, and inflammation regulation, suggesting a tumor-promoting immunosuppressive microenvironment. *ALOX5, NCF4*, and *CYBB* (likelyincluded as part of the NADPH oxidase complex) are involved in reactive oxygen species (ROS) production, linking oxidative stress to tumor progression and resistance to cell death.

**Figure 2.**
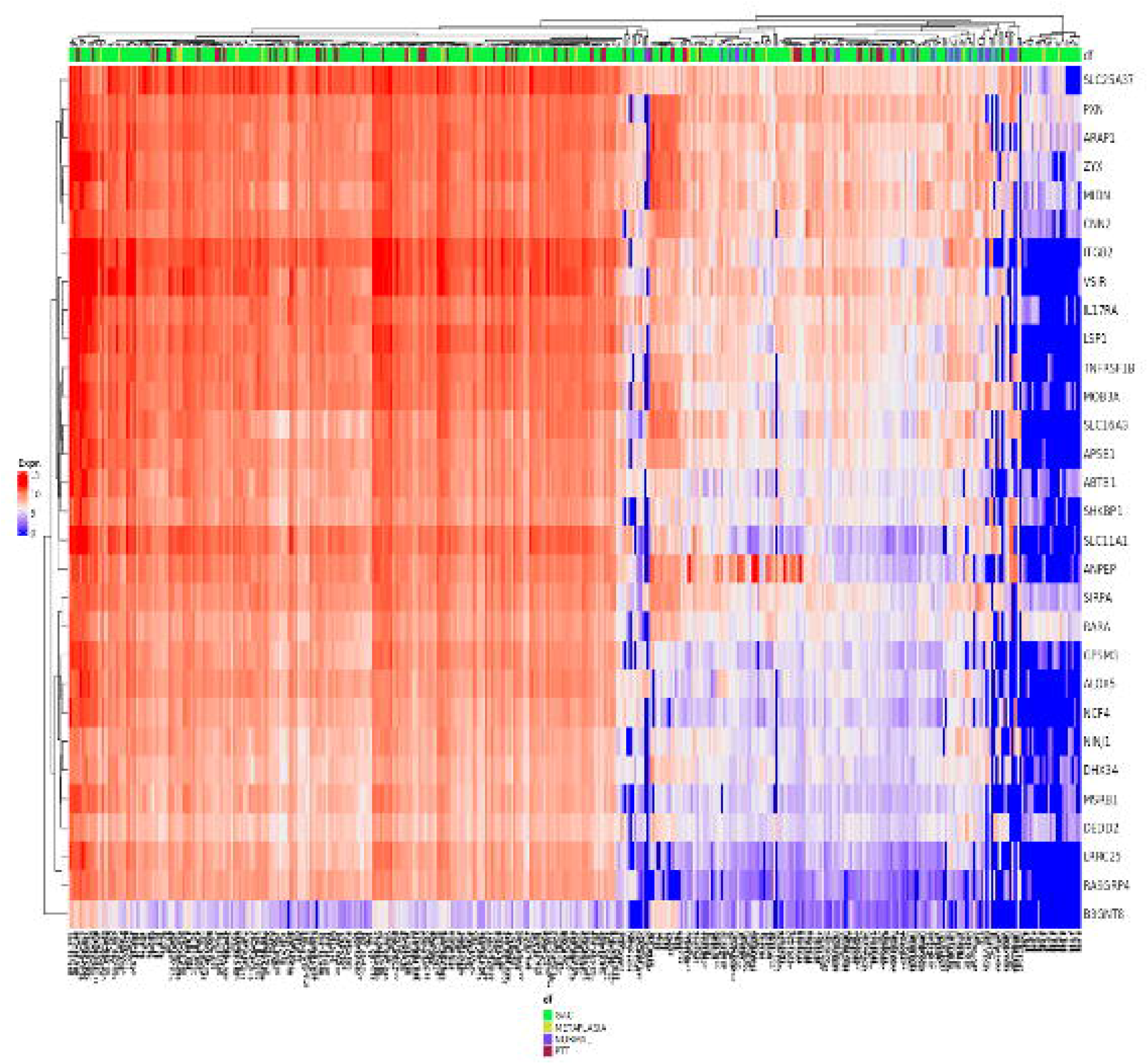
Heatmap of differentially expressed genes in gastric cancer samples.

Several solute carrier genes, such as *SLC25A37, SLC16A3*, and *SLC11A1*, highlight metabolic reprogramming, particularly in mitochondrial iron transport and lactate export, essential for sustaining cancer cell proliferation.

Transcriptional regulators like *SPI1* and *RARA*, along with chromatin remodelers like *SMARCB1*, underscore the role of epigenetic and transcriptional dysregulation in GC pathogenesis.

Additionally, genes such as *NINJ1* and *DEDD2*, associated with cell death pathways, may indicate adaptations that modulate apoptosis and pyroptosis.

These findings highlight a coordinated upregulation of genes involved in immune suppression, oxidative stress, metabolic adaptation, and cell survival, all of which contribute to gastric cancer progression and therapy resistance.

### Correlation Analysis

The correlation analysis involving the gene *ANPEP* revealed a set of genes whose expression patterns were positively associated with *ANPEP* across multiple tumor samples (Figure 3). These associations suggest potential biological interactions and functional relevance within the tumor microenvironment, particularly in the context of immune regulation, cell migration, and tumor progression.

**Figure 3.**
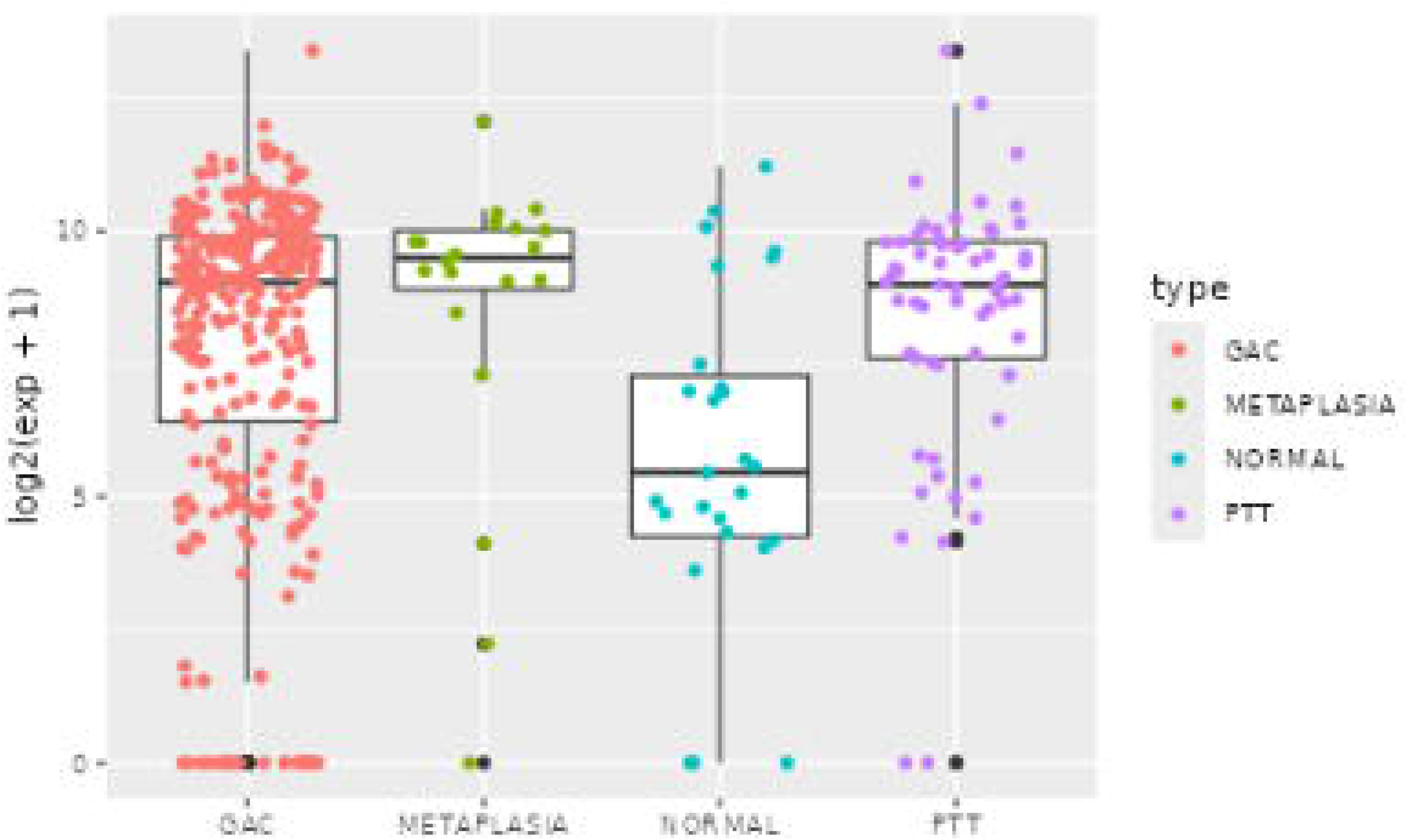
Correlation analysis between ANPEP expression and DE genes with more than 10 reads (normalized expression) in at least 20 samples.

Several genes exhibited strong positive correlations with *ANPEP*,indicating that they may participate in shared biological pathways. Notably, *PXN* (*Paxillin*), a focal adhesion protein involved in cytoskeletal dynamics and cell motility, showed concordant expression with *ANPEP*, suggesting a possible role in enhancing migratory and invasive behavior of tumor cells. This aligns with research showing that cytoskeletal remodeling is critical for tumor cell invasion and metastasis (ASEERVATHAM *et al*., 2020).

Similarly, *TNFRSF1B*, a receptor of the TNF superfamily involved in inflammation and cell survival, was highly expressed alongside *ANPEP*, highlighting a potential link between *ANPEP* and inflammatory signaling within the tumor microenvironment, which can promote tumor progression through immune modulation.

*SIRPA*, an inhibitory immune receptor, was also positively correlated with *ANPEP*. This gene is known to inhibit phagocytosis by macrophages, potentially aiding tumor cells in escaping immune surveillance (MHAIDLY *et al*., 2020).

This association implies a functional connection between *ANPEP* expression and the regulation of innate immune responses, including macrophage activation and immune evasion. *VSIR* (also known as *VISTA*), a negative immune checkpoint regulator, showed strong co-expression with *ANPEP*, suggesting a possible synergistic role in suppressing adaptive immune responses and promoting immune tolerance within the tumor, consistent with its role in creating an immunosuppressive microenvironment (HUANG *et al*., 2024).

Additional positively correlated genes included *ZYX* (*Zyxin*), involved in actin cytoskeleton organization; *ARAP1*, which regulates cytoskeletal remodeling via *ARF* and *Rho* signaling pathways; and *CNN2* (*Calponin 2*), a protein associated with cell contraction and motility. Their co-expression with *ANPEP* reinforces its putative involvement in cytoskeletal regulation and tissue remodeling processes that are characteristic of invasive tumors, as cytoskeletal dynamics are crucial for tumor cell motility and metastasis (GUO *et al*., 2023).

The integrin gene *ITGB2*, known for its role in leukocyte adhesion and signaling, was also highly correlated with *ANPEP*, suggesting possible coordination in modulating leukocyte infiltration and intercellular adhesion, which can influence immune cell recruitment to the tumor site (NEO *et al*., 2020). Similarly, *SLC25A37*, a mitochondrial iron transporter, and *LSP1*, which modulates leukocyte motility, were positively associated with *ANPEP*, pointing to a metabolic and immunological convergence possibly related to increased cellular turnover, as metabolic reprogramming is a hallmark of tumor progression (LIU *et al*., 2024).

These findings support a model in which elevated *ANPEP* expression is part of a coordinated transcriptional program involving genes that promote immune modulation, cell migration, and metabolic adaptation, hallmarks of tumor progression and immune evasion. These data provide a molecular rationale for further investigating *ANPEP* as both a biomarker and a functional mediator in tumor biology.

## DISCUSSION

The present study provides compelling evidence for the multifaceted role of *ANPEP* in GC progression, highlighting its potential as both a biomarker and therapeutic target. The observed upregulation of *ANPEP* across gastric GC, PTT, and M compared to N tissues (p < 0.01) aligns with previous findings that link *ANPEP* overexpression to early mucosal transformation and malignant progression in GC (HOFT *et al*., 2024).

The heatmap analysis revealed the upregulation of a subset of genes, comprising immune-related genes (e.g., *ITGB2, VSIR, IL17RA, TNFRSF1B*) and inflammation/oxidative stress genes (e.g., *ALOX5, NCF4, NINJ1, DHX34*) - further elucidates *ANPEP*”s contribution to an immunosuppressive tumor microenvironment (TME) (ZHANG *et al*., 2017). The upregulation of *VSIR* (*VISTA*), a negative immune checkpoint regulator (LEMERCIER et al., 2014), supports the notion of a tumor phenotype with reduced immune surveillance (HUANG *et al*., 2024). This finding is corroborated by studies showing that downregulation of immune checkpoint pathways enhances tumor immune evasion, a hallmark of advanced GC (NEO *et al*., 2020).

*ALOX5, NCF4*, and *CYBB*, components of the NADPH oxidase complex, are integral to ROS generation. Increased ROS levels can promote DNA damage, genomic instability, and activate signaling pathways that enhance tumor cell proliferation and survival. Specifically, *NOX4*, a member of the NADPH oxidase family, has been shown to regulate GC cell proliferation and apoptosis through the *GLI1* pathway, highlighting the role of ROS in GC pathogenesis (TANG et al., 2018).

Transcription factors like *SPI1* and *RARA*, along with chromatin remodelers such as *SMARCB1*, are associated with transcriptional dysregulation. *SPI1* has been implicated in promoting GC progression by activating the IL6/JAK2/STAT3 signaling pathway (HOU et al., 2022).

The correlation analysis reinforces *ANPEP*”s involvement in a coordinated transcriptional network. Positive correlations with genes like *PXN, ZYX*, and *CNN2* underscore its role in cytoskeletal dynamics and cell motility, aligning with evidence that cytoskeletal remodeling drives GC metastasis (GUO *et al*., 2023). The co-expression of *ANPEP* with *TNFRSF1B* and *VSIR* highlights its immunomodulatory potential, potentially exacerbating inflammation and immune tolerance, as supported by studies on TNF signaling in GC (OSHIMA *et al.*, 2014).

These findings are consistent with the hypothesis that *ANPEP* contributes to a tumor-promoting TME through immune evasion and metabolic adaptation and poses *ANPEP* as a central player in GC biology, bridging molecular, and immunological factors.

The upregulation of *ANPEP* in pre-malignant and malignant states, coupled with its association with an immunosuppressive and plastic TME, provides a molecular basis for its potential as a prognostic biomarker and therapeutic target. Targeted therapies, such as *ANPEP* inhibitors could be explored, particularly in combination with immune checkpoint inhibitors, to overcome the immunosuppressive barriers identified.

## CONCLUSION

This study demonstrates that *ANPEP* is consistently upregulated from early pre-malignant stages through to advanced gastric cancer, independent of conventional clinicopathological parameters. Its expression is tightly linked with a network of genes involved in immune suppression, oxidative stress, metabolic reprogramming, cytoskeletal remodeling, and cell survival pathways. These coordinated transcriptional changes suggest that *ANPEP* plays a central role in shaping an immunosuppressive, metabolically adaptive, and invasion-prone tumor microenvironment that facilitates gastric cancer progression and immune evasion. The strong associations between *ANPEP* and key immune checkpoint regulators (e.g., *VSIR*), inflammatory mediators (e.g., *TNFRSF1B*), and cytoskeletal components (e.g., *PXN, CNN2*) further highlight its multifaceted contribution to tumor biology. These findings position *ANPEP* as a promising biomarker for early detection and a potential therapeutic target, particularly in strategies aiming to disrupt tumor immune escape and enhance treatment efficacy in gastric cancer. Importantly, the markedly low expression of *ANPEP* in normal gastric tissues further supports its potential as a therapeutic target, offering the advantage of high specificity for transformed cells while minimizing off-target cytotoxicity.

## ACKNOWLEDGMENTS

The authors are grateful to the Brazilian Agencies CNPq, CAPES and Federal University of Pará (PROPESP and FADESP) for fellowships and financial support.

## DATA AVAILABILITY STATEMENT

All relevant data is contained within the article: The original contributions presented in the study are included in the article, further inquiries can be directed to the corresponding author.

## ETHICS STATEMENT

The studies involving humans were approved by Ethics Committee for Research of the João de Barros Barreto University Hospital (CAEE - 47580121.9.0000.5634). The studies were conducted in accordance with the local legislation and institutional requirements. The participants provided their written informed consent to participate in this study. Written informed consent was obtained from the individual(s) for the publication of any potentially identifiable images or data included in this article.

## AUTHOR CONTRIBUTIONS

Conceptualization, PPA and TMTA; sample collection, WFB, GI and AKMA; methodology, SMMC, JMCS, RMSM, EBT, DP, VCSS, DSA, RFS, AV, LM; software and data curation, FCM; writing-original draft preparation, TMTA, MSNR and BFRR; writing-review and editing, AKRS, LECM, RRB, PPA; supervision, PPA. All authors have read and agreed to the published version of the manuscript.

